# Vacuolar transporter Mnr2 safeguards mitochondrial integrity in aged cells

**DOI:** 10.1101/2020.07.23.217471

**Authors:** Md. Hashim Reza, Rajesh Patkar, Kaustuv Sanyal

**Author notes:** Biological and Life Sciences Division, School of Arts and Sciences, Ahmedabad University, Ahmedabad 380 009, Gujarat, India (Rajesh Patkar).

## Abstract

Aging is associated with altered mitochondrial function. Mitochondrial function is dependent on the magnesium (Mg^+2^) ion flux. The molecular mechanism underlying Mg^+2^ homeostasis, especially during aging has not been well understood. We previously demonstrated that the absence of a vacuolar ion transporter Mnr2 accelerates cell death in the older part of the colony in *Magnaporthe oryzae* presumably due to an altered Mg^+2^ homeostasis. Localization of Mnr2 as dynamic puncta at the vacuolar membrane especially in the older *Magnaporthe* cells further suggests its association with aged cells. Interestingly, such vacuolar Mnr2 puncta colocalized with the filamentous mitochondria in the aged cells. Further, we show that aged *mnr2*Δ null cells displayed loss of integrity of mitochondria and vacuoles. Remarkably, exogenously added Mg^+2^ restored the mitochondrial structure as well as improved the lifespan of *mnr2*Δ null cells. Thus, we uncover a mechanism of maintenance of mitochondrial integrity and function by the ion transporter Mnr2-based Mg^+2^ homeostasis during aging.

## Introduction

Aging, a time-dependent decline in biological fitness, is characterized by a progressive accumulation of impaired cellular function leading to an increased susceptibility to death. Repeated damage and metabolic perturbations contribute to aging but are kept under control by a complex network of maintenance and repair functions. Over the years, various model systems including worms, flies, mice, and budding yeast have been employed to study the process of aging (Gershon and Gershon 2000; Brandt and Vilcinskas 2013; Tissenbaum 2015; Folgueras *et al.* 2018). Aging in yeast can be assayed by measuring replicative lifespan (RLS) that is measured by the number of times an individual cell divides, or chronological lifespan (CLS) which measures the length of time that a non-dividing cell survives (Longo *et al.* 2012; Carmona-Gutierrez and Buttner 2014). Several factors which contribute to aging include genome instability, telomere attrition, epigenetic alterations, impaired protein homeostasis, deregulated nutrient sensing, cellular senescence, altered intercellular communication, stem cell exhaustion, and mitochondrial dysfunction (Lopez-Otin *et al.* 2013; Lopez-Otin *et al.* 2016).

In yeast, mitochondrial function is important for both replicative and chronological lifespan (Longo *et al.* 2012). While mitochondria influence different aspects of cellular senescence, they also are a target of the process, and are thus central to the process of aging (Shigenaga *et al.* 1994; Kirkwood 2005; Nisoli *et al.* 2005; Rea *et al.* 2007). several studies have also highlighted the role of acidic vacuoles in lifespan extension, either through regulating mitochondrial structure and function or through autophagy (Melendez *et al.* 2003; Hughes and Gottschling 2012; Ruckenstuhl *et al.* 2014). Autophagy is essential during chronological aging for the recycling of resources. Further, vacuolar fusion during nutrient restriction has been shown to extend lifespan in yeast (Tsuchiyama and Kennedy 2012). Vacuoles have a role not only in degradation and autophagy, but also in ion homeostasis via storage of various metabolites, vacuolar protein sorting, and detoxification. The degradative capacity of vacuoles is largely dependent on the acidic milieu, which is maintained by the activity of V-ATPase (Kane 2006), which in turn is regulated by various proton antiporters of metal ions and amino acids. The acidity of the vacuoles declines with age. A recent study reports a previously unknown correlation between the pH maintenance inside the vacuole, and its function in prolonging mitochondrial integrity in yeast during replicative aging (Hughes and Gottschling 2012).

Mitochondria need constant communication and exchange of small molecules with other organelles as it serves as the hub for signaling required for development, differentiation, and cell death. Membrane contact sites (MCSs) have evolved for metabolic communication between various organelles within a cell without fusing to represent another elegant means of intra-cellular communication. MCSs are pivotal for vesicular independent transport to facilitate exchange of ions, metabolites, and lipids across short distances between membranes. While ER–mitochondrial contacts are long known, contacts between the endolysosomal system and mitochondria only recently came into the limelight (Elbaz-Alon *et al.* 2014).

In a screen to identify Vps39-interacting proteins, Mnr2 was identified (Elbaz-Alon *et al.* 2014). As a member of the Cobalt Resistance A (CorA) Mg^+2^ transporter superfamily, Mnr2 is a potential component of vacuole and mitochondria patch (vCLAMP) in yeast. While transporters at the plasma membrane help in the uptake of nutrients from the external environment into the cytosol, those on the vacuolar membrane contribute to homeostasis of molecular building blocks and ions, especially under stress and/ or nutrient depletion conditions. Both vacuoles and mitochondria are major storage organelles for magnesium (Mg^+2^) ions (Kubota *et al.* 2005; Pisat *et al.* 2009). Magnesium ions are required for stabilization of proteins and nucleic acids, regulation of function of ion channels, and cofactors in mediating enzymatic reactions in a cell (Wolf and Cittadini 2003). The normal physiological functioning of a cell is therefore dependent on proper homeostasis of the intracellular Mg^+2^. Dysregulation of mitochondrial Mg^+2^ homeostasis disrupts ATP production plausibly due to an alteration in the energy metabolism and morphology of the organelle (Yamanaka *et al.* 2016). Deficiency in the intracellular Mg^+2^ concentration is also associated with neuronal and age-related diseases, including accelerated cellular senescence (Longstreth *et al.* 2002; Mario *et al.* 2011), whereas excess cytoplasmic Mg^+2^ can cause toxic effects such as inactivation of key enzymes involved in carbon fixation in *Spinacia oleracea L.* (Wu *et al.* 1991).

*M. oryzae* is an economically important filamentous ascomycete plant pathogen that causes devastating blast disease on various cereal crops including rice and wheat (Wilson and Talbot 2009; Islam *et al.* 2016). It is considered as a model system to study fungal pathogenesis and host-pathogen interaction. We have previously shown that the demand for intracellular Mg^2+^ increases during developmental progression, from vegetative hyphal growth to sporulation and germination to appressorium formation, in *Magnaporthe oryzae* (Reza *et al.* 2016). Further, unlike the plasma membrane Mg^+2^ transporter Alr2, which is essential for viability and pathogenesis, loss of Mnr2 function led to a significant cell death in the older part of the colony of *M. oryzae* (Reza *et al.* 2016). A similar reduction in viability in the older part of the vegetative culture was observed upon loss of Ech1, the mitochondrial beta-oxidation enzyme, function in *M. oryzae* (Patkar *et al.* 2012; Reza *et al.* 2016). However, the mechanistic insights underlying cellular fitness during aging and any cross talk between organellar function contributing towards longevity in *M. oryzae* remain unclear thus far. in the present study, we describe the role of the vacuolar Mg^+2^ transporter, Mnr2, in organellar integrity and chronological lifespan in *M. oryzae*. We show the age-specific temporal expression and dynamics of Mnr2, and provide evidence that Mnr2 plays a key role for transporter-driven Mg^+2^ homeostasis in maintaining the integrity and function of mitochondria and to facilitate survival of chronologically older cells.

## Results

### Mnr2 is required for sustained lifespan and is expressed predominantly in aged cells of *Magnaporthe oryzae*

Filamentous fungi, where a daughter cell remains attached to the parent, grow as a radial mycelial network emanating from the point of inoculation that is the center of a colony. Thus, the cells at the center are the oldest while those at the hyphal tips along the colony periphery are the youngest, forming an age gradient across the fungal colony (Figure 1A). We previously characterized two CorA magnesium transporters, Alr2 and Mnr2, in the rice-blast fungus *M. oryzae* having a role in fungal development and progression of infection cycle (Reza *et al.* 2016). Mnr2 related proteins are absent in plants and metazoans. Here, we identified the putative homologs of Mnr2, having conserved CorA domain and WGMN sequence, in species across Ascomycota and Basidiomycota using *in silico* predictions (Supplementary Figure S1). The length of the CorA domain towards the C-terminus is significantly conserved across species in both the phyla (Supplementary Figure S1).

**Figure 1:**
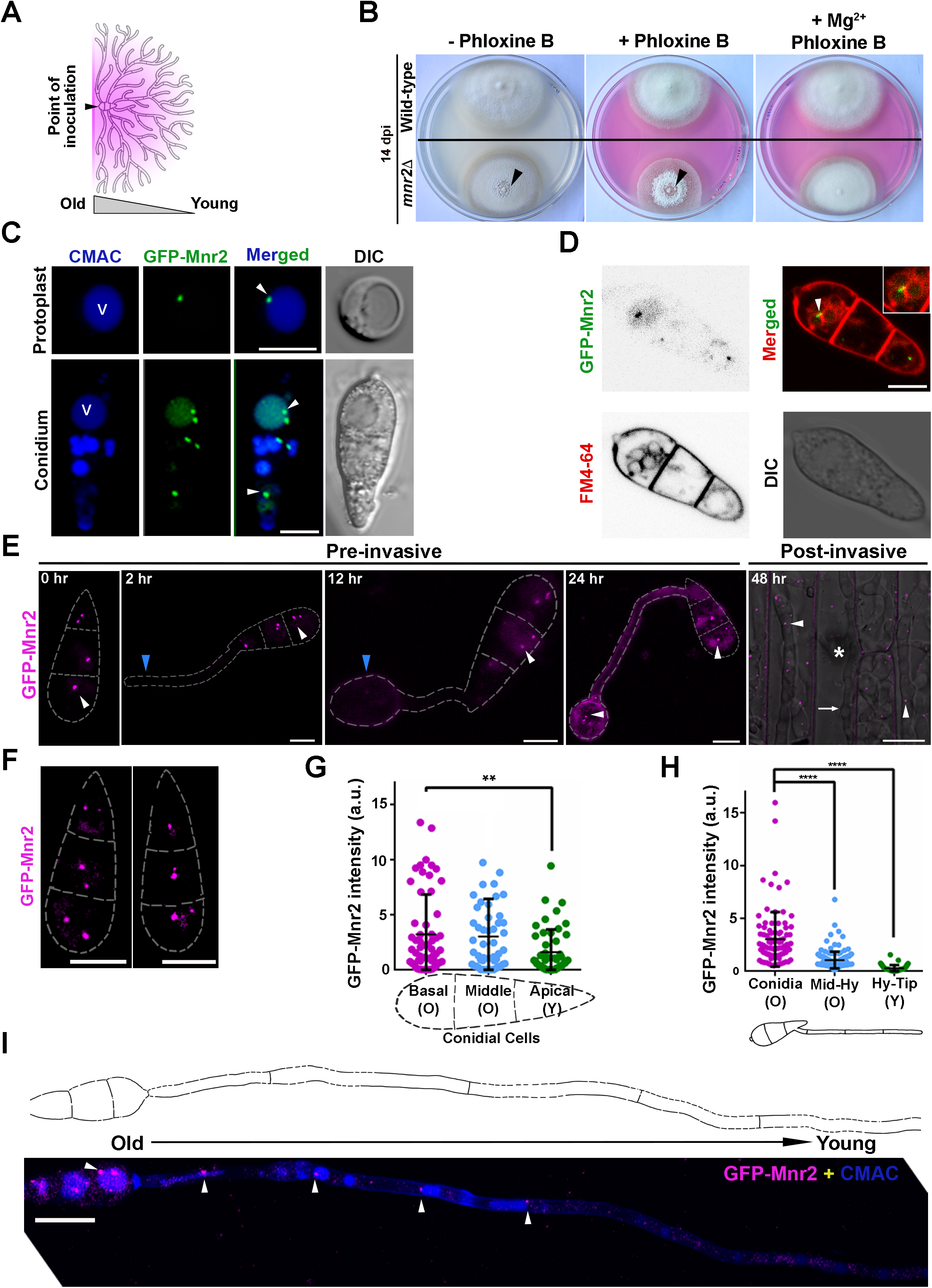
Mnr2 is required for survival fitness in chronologically aged cells and is expressed specifically in the old cells of *M. oryzae.* (A) A schematic depicting the relative age of cells in a radial filamentous fungal colony. The triangle marks the older (point of inoculation) and younger (periphery of the colony) parts of the vegetative colony. (B) Growth of the wild-type and *mnr2*Δ *M. oryzae* colonies, grown on YEGA, YEGA with phloxine B, or YEGA supplemented with Mg^+2^ and phloxine B, 14 days post inoculation (dpi). Arrowheads mark the loss of viability at the central, aged part of the colony. The data represents observation from three independent experiments. (C) Laser-scanning confocal micrographs depicting the subcellular localization of GFP-Mnr2 in protoplasts (n = 23) and conidia (n = 66). Both protoplasts and conidia were stained with CMAC to aid visualization of vacuoles, marked as V. Arrowheads mark the GFP-Mnr2 puncta. Scale bar, 5 μm. (D) A representative maximum-intensity projection image of the wild-type conidium showing GFP-Mnr2 localization at the vacuolar membrane marked by FM4-64 staining. Inset, a magnified region depicted by an arrowhead. Scale bar, 5 μm. (E) Expression of GFP-Mnr2 during pre-invasive, on glass coverslips, and post-invasive, on rice leaf sheath, stages of pathogenic development in *M. oryzae* - conidia (0 h), conidial germination (2 to 3 h), young appressorium (12 h), mature appressorium (24 h), and invasive hyphae (48 h). White arrowheads mark the GFP-Mnr2 puncta at the vacuolar periphery, whereas blue arrowheads depict the young cells of germ tube and young appressorium with no GFP-Mnr2 expression. Asterisk shows the appressorium that penetrated and elaborated the invasive hyphae (white arrow) within the rice leaf sheath cell. Scale bar, 5 μm (pre-invasive) and 10 μm (post-invasive). (F) Representative maximum-intensity projection images showing GFP-Mnr2 expression in all the three cells of conidia. Scale bar, 5 μm. (G) A scatter plot depicting the differential intensity of the GFP-Mnr2 puncta in the three (basal, middle, and apical) cells of conidia. A schematic is shown as a part of the *x*-axis label to mark the three cells of a conidium and their corresponding age. O, older cell; Y, Young cell. Levels indicate normalized GFP-Mnr2 intensity for each analyzed cluster. Data represents mean ± SEM from three independent experiments and calculated using one-way ANOVA followed by Dunnett’s multiple comparison test. n = 51 to 58 conidia per replicate. ** indicates *P* = 0.05. (H) A scatter plot showing the differential intensity of the GFP-Mnr2 puncta in different cell compartments of a vegetative hypha. A schematic is shown as a part of the *x*-axis label to depict the hyphal cellular compartments of different chronological age. O, older cells; Y, younger cells; Mid-Hy, mid hyphal cells; Hy-Tip, hyphal tip cells. Levels indicate normalized GFP-Mnr2 intensity for each analyzed cluster. Data represents mean ± SEM from three independent experiments and calculated using one-way ANOVA followed by Dunnett’s multiple comparison test. n = 52 to 162 hyphae per replicate. **** indicates *P* = 0.05. (I) Expression of GFP-Mnr2 in a chronological age-dependent manner in a vegetative hypha. A schematic shows an outline of the cellular compartments in the vegetative hypha for which corresponding laser-scanning confocal micrographs are shown in the panels below. The direction of arrow indicates the age gradient from the old cells in the conidium to young cells towards the tip of the growing hypha. Arrowheads show colocalization of GFP-Mnr2 punctae, pseudo colored to magenta, with CMAC-stained vacuoles in older cells. Scale bar, 10 μm.

While the membrane Mg^+2^ transporter Alr2 is essential for viability, the vacuolar counterpart Mnr2 plays a role in fungal development and pathogenesis. The loss of Mnr2 function leads to premature cell death in the fungal culture grown on a laboratory medium (Reza *et al.* 2016). Upon staining with the vital dye phloxine B, here we observed an accelerated cell death in the central part of the 13- to 14-day aged vegetative colony of the *mnr2*Δ null mutant (Figure 1B). Thus, upon loss of Mnr2 function, the chronologically aged cells at the central part of the culture, lost viability (Figure 1B). As shown earlier (Reza *et al.* 2016), exogenously added Mg^+2^ significantly restored viability in the *mnr2*Δ culture (Figure 1B), indicating a likely role for Mnr2-based Mg^+2^ homeostasis in the survival fitness of chronologically aged cells in *Magnaporthe.*

To understand the subcellular localization of Mnr2 in *M. oryzae*, we first functionally expressed an N-terminally GFP-tagged Mnr2 protein in the wild-type background (hereinafter referred to as GFP-Mnr2) (Supplementary Figure S2). Confocal microscopy revealed that GFP-Mnr2 was localized as one or two discrete puncta at the vacuoles in all the three conidial cells and protoplasts (Figure 1C). The localization of GFP-Mnr2 puncta at the vacuolar membrane was further confirmed by co-staining of vacuolar membrane with FM4-64 dye (Figure 1D). In contrast to uniform membrane localization, such spatially confined focal structures, representing a unique cellular niche, are commonly observed in organelle contact sites like endoplasmic reticulum mitochondrial encounter sites (ERMES), vCLAMP, and nuclear-vacuole junctions (NVJ) (Elbaz-Alon *et al.* 2014).

Next, we studied the expression of GFP-Mnr2 at different stages of fungal development. Here, we followed the expression of GFP-Mnr2 during both pre-invasive (three-celled conidium, 0 hpi; conidial germination, 2 hpi; appressorial development, 12 & 24 hpi) and post-invasive (invasive hyphae, 48 hpi) pathogenic development in *M. oryzae.* Intriguingly, not all the fungal cells, irrespective of the developmental stage, showed GFP-Mnr2 puncta. For instance, while all the three cells of a conidia displayed punctate localization of GFP-Mnr2, germinating conidia at 2 to 3 hpi showed puncta particularly in the conidial cells but not in the newly emerged germ tubes (blue arrowheads; Figure 1E). Similarly, incipient/ immature appressoria (12 hpi) rarely displayed GFP-Mnr2 puncta (blue arrowheads; Figure 1E). Importantly, as the appressoria matured (24 hpi), the expression of GFP-Mnr2 was evident therein (white arrowhead; Figure 1E). Similarly, the invasive hyphal cell compartments (48 hpi) revealed the expression of GFP-Mnr2 puncta at the vacuolar periphery.

At a close examination, there seemed to be differences in GFP-Mnr2 expression across the three cells of the conidium. In *M. oryzae*, conidia formation is marked with three sequential rounds of mitotic events (Shah *et al.* 2019). In the chronological order, the basal cell is older than the apical/ terminal cell in a conidium. We quantified the intensity of GFP-Mnr2 in all the three cells and found that indeed the expression of GFP-Mnr2 was significantly higher in older (basal and middle) cells when compared to the young (apical) cell (Figure 1F & 1G; *P* = 0.05). To further explore the possibility that Mnr2 express in an age-dependent manner, we studied the expression of GFP-Mnr2 in vegetative hyphae grown from germinating conidia. Hyphal growth in filamentous fungi occurs by extension at the tips in a monopolar fashion, followed by cell division giving rise to apical/ tip (young) and subapical (old) cells. Consistent with the aforementioned pattern, GFP-Mnr2 did not express in the apical/ tip (young) cell of the vegetative hyphae in *M. oryzae* (Figure 1H & 1I). Rather, interestingly, GFP-Mnr2 puncta were seen with gradually increasing intensity, in a chronological age-dependent manner, in aged cells along the vegetative hyphae and conidia (Figure 1H & 1I). While the old cells showed significantly higher GFP-Mnr2 expression, that in the young/ tip cells was almost negligible (Figure 1H & 1I; *P* = 0.05).

To verify our observation that Mnr2 was expressed predominantly in the aged cells, we expressed GFP under the native *MNR2* promoter, and observed the intensity of the reporter protein in the vegetative hyphae in *M. oryzae.* As a control, we separately studied the expression of GFP driven by a constitutive *MPG1* promoter. We found that the *MNR2*-promoter-driven GFP was significantly expressed in the chronologically older conidia and hyphal cells away from the tips when compared to the young apical/ tip cells (Supplementary Figure S3A & S3B; *P* <0.005). Earlier studies highlighting the role of acidic vacuoles and of vacuolar fusion during nutrient restriction towards lifespan extension in yeast (Tsuchiyama and Kennedy 2012), intrigued us to look at vacuolar functions in the *mnr2*Δ mutant.

### Mnr2 plays a crucial role in maintaining the vacuolar integrity and function in *M. oryzae*

The vCLAMP proteins Vps39 and Ypt7 are involved in maintaining vacuolar integrity in the budding yeast (Balderhaar *et al.* 2010). Since Mnr2 is a Vps39-mediated vCLAMP resident protein, we asked whether Mnr2 had any role in maintaining the vacuolar integrity in *M. oryzae.* Indeed, approximately 55% of the *mnr2*Δ conidia possessed multiple and small spherical vacuoles as against few yet larger vacuoles in the wild-type conidia (Figure 2A). Since, vacuolar function is dependent on the acidic pH of the lumen, we carried out quinacrine staining of the wild-type and *mnr2*Δ mycelia grown under nutrient starvation conditions. We found that the wild-type had a greater number of quinacrine-stained highly acidic vacuoles when compared to those in the *mnr2*Δ, wherein mostly the cytoplasm was stained with the dye (Figure 2B & 2C; *P* <0.005). The V-ATPase mutants of yeast are shown to be sensitive to oxidative stress (Li and Kane 2009). Given the defect in pH homeostasis, as revealed by quinacrine staining in the *mnr2*Δ mutant, we studied sensitivity of the mutant towards oxidative stress. Indeed, the *mnr2*Δ mutant was more sensitive to H2O2 when compared to the wild-type (Figure 2D).

**Figure 2:**
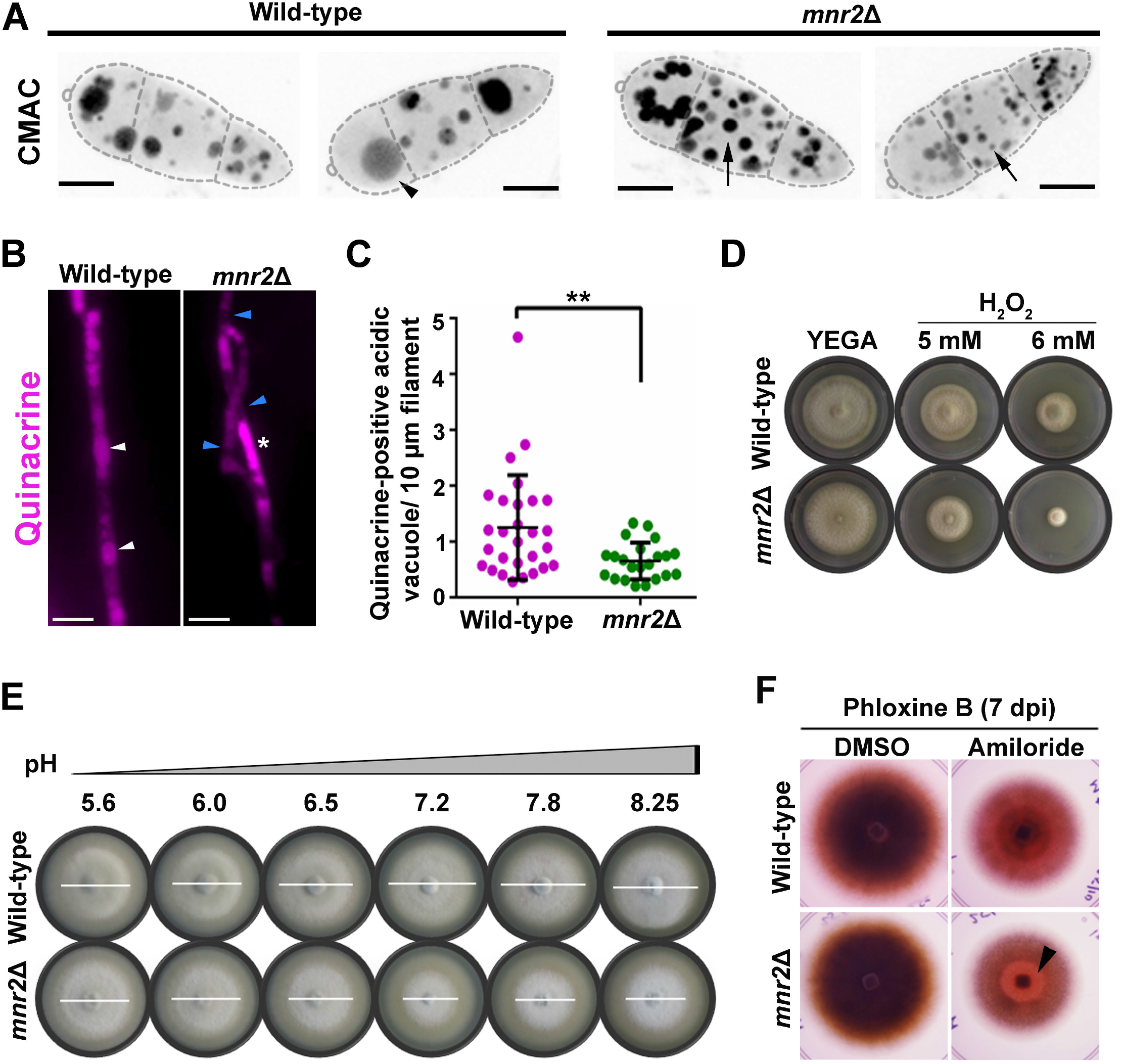
Mnr2 plays a crucial role in vacuolar integrity and function. (A) Laser-scanning confocal micrographs depicting CMAC-stained vacuoles with different sizes and numbers in the wild-type (n = 40) and *mnr2*Δ (n = 36) strains of M. oryzae. Arrowhead shows larger vacuoles in the wild-type, while arrows mark the smaller vacuoles in the *mnr2*Δ mutant. Scale bar, 5 μm. (B) Vacuolar pH of the vegetative hyphae, under nutrient starvation condition, assessed by staining with quinacrine. White and blue arrowheads mark quinacrine-stained (low pH) and unstained (high pH) vacuoles, respectively, in the wild-type and *mnr2Δ.* Asterisk marks quinacrine-stained cytoplasm in the *mnr2*Δ hyphal cells. Scale bar, 10 μm. (C) A scatter plot showing the quantitative analysis of the quinacrine-stained vacuoles per 10 μm stretch of hyphae. n = 28 (wild-type) and 22 (*mnr2Δ).* ** denotes P = 0.0033 (unpaired t-test). (D) Vegetative growth of the wild-type and *mnr2*Δ on YEGA medium with or without the indicated concentrations of H2O2, at 10 dpi. (E) Vegetative growth of the wild-type and *mnr2*Δ on YEGA medium with indicated pH at 5 dpi. White lines mark the diameters of the respective fungal colonies. (F) Effect of amiloride HCl-mediated inhibition of Mg+2/H+ exchanger on longevity of the aged cells, assessed by staining with the vital dye phloxine B, of the wild-type and *mnr2*Δ cultures. Black arrowhead depicts accelerated cell death on 7 dpi at the centre, marking the aged cells of the *mnr2*Δ colony.

Next, to study whether the altered vacuolar pH also affected the autophagic turnover of mitochondria, we deleted *MNR2* in the Atp1-GFP-expressing strain of *M. oryzae* (Patkar *et al.* 2012) (Supplementary Figure S4). The extent of mitophagy was assessed in the Atp1-GFP or Atp1-GFP/ *mnr2*Δ strain grown under nitrogen starvation conditions. We found that overall mitophagy, read as GFP signal intensities in the vacuolar lumen, was impaired in the *mnr2*Δ strain when compared with the wild-type and was similar to wild-type treated with PMSF (Supplementary Figure S5A). Similarly, we also tested laccase activity as a measure of vacuolar function in both wild-type and *mnr2*Δ strains. While the specific laccase activity in the wild-type was over 6 U/ mg at 48 hours post inoculation (hpi) that in the *mnr2*Δ mutant was drastically reduced to 0.028 U /mg (Supplementary Figure S5B).

Non-acidic vacuole mutants of yeast show inability to grow at alkaline pH (Kane 2006). Having shown a defective vacuolar pH in the *mnr2*Δ mutant, we then asked whether the *mnr2*Δ mutant had developed sensitivity to the pH of the extracellular medium. We found that the *mnr2*Δ colony diameter was smaller at neutral and alkaline pH when compared with that of the wild-type (Figure 2E). The correct vacuolar pH and Mg^+2^ ion homeostasis is likely achieved by a concerted activity of the V-ATPase, Mnr2, and Mg^2+^/ H^+^ exchanger (Pisat *et al.* 2009). Alterations in the function of any one of these transporters may affect the activity of the other. We surmised that inhibition of the exchanger, which functions opposite to Mnr2 and V-ATPase, might restore the wild-type-like phenotype in the *mnr2*Δ mutant. On the contrary, we found that the treatment of the *mnr2*Δ with amiloride HCl further accelerated the cell death in the mutant, where an autolysis was seen as early as 7 days as compared with 14 days in *mnr2*Δ without addition of inhibitor (Figure 2F). This suggests that a fine balance between the functions of vacuolar Mg^2+^ and H^+^ transporters is crucial for ion homeostasis and sustained chronological lifespan in *M. oryzae.*

### Vacuolar transporter Mnr2 colocalizes with mitochondria in the old cells of *M. oryzae*

Having shown GFP-Mnr2 to be localized as a clustered punctum and with its expression being higher in aged cells of *M. oryzae*, we next asked whether GFP-Mnr2 showed dynamic sub-cellular localization. Live-cell imaging revealed that GFP-Mnr2 puncta were highly dynamic and moved along the vacuolar periphery (Figure 3A; Supplementary Movie 1). We further found that the number of puncta per cell was dynamic and often associated with the change in the punctum size (Figure 3B). It is likely that the different sizes of GFP-Mnr2 puncta resulted from differential clustering of more than one punctum/ molecule of Mnr2.

**Figure 3:**
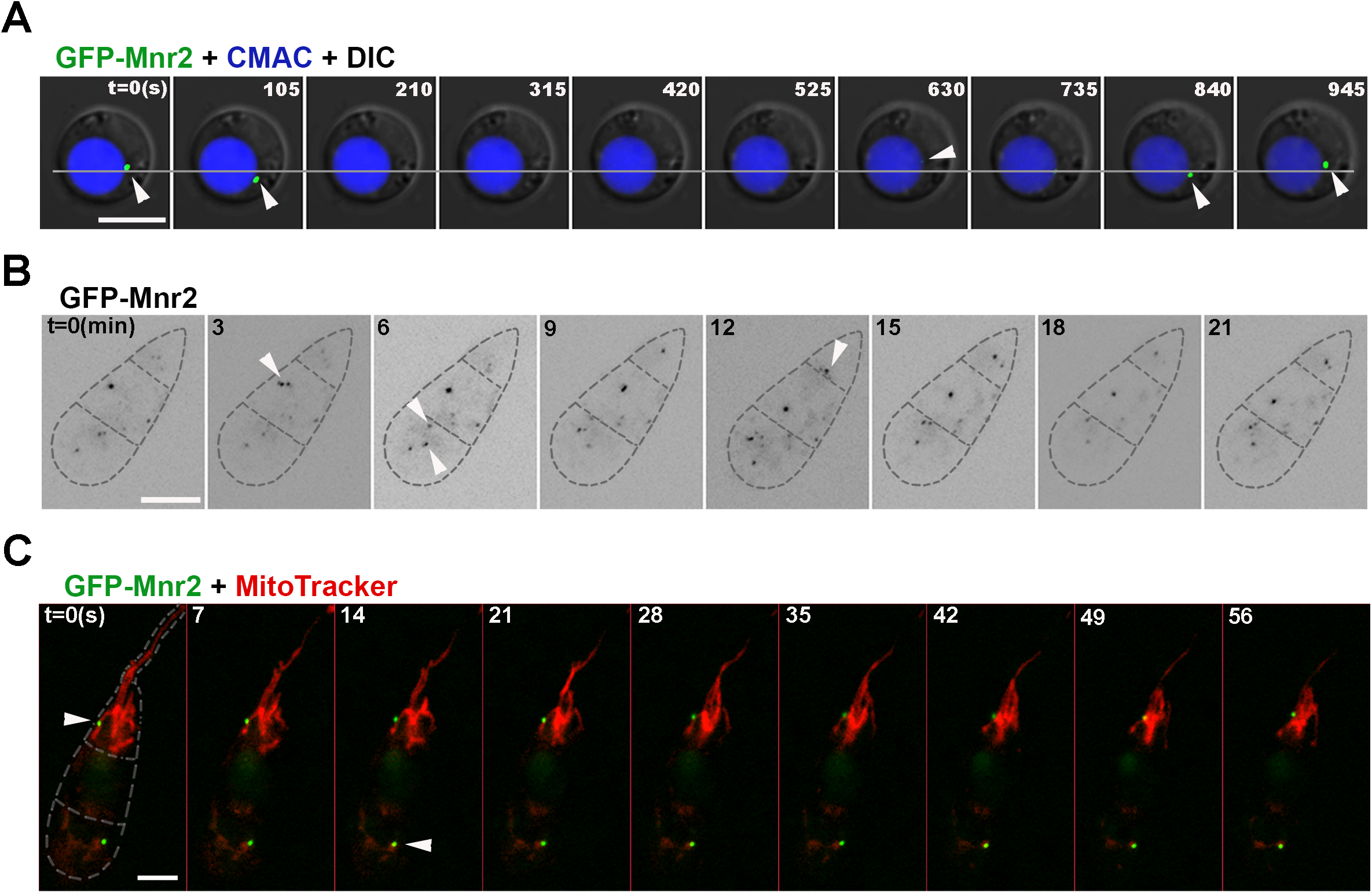
Dynamic vacuolar membrane transporter protein Mnr2 colocalizes with filamentous mitochondria in aged cells. (A) A representative montage of time-lapse images showing dynamic movement of a GFP-Mnr2 puncta in a protoplast. Arrowheads depict the GFP-Mnr2 puncta along the periphery of the CMAC-stained vacuole. A line, marking the initial position of a GFP-Mnr2 puncta in the first frame, is drawn across the montage to indicate the shift in the position of the same puncta at subsequent time points. Numbers indicate time in seconds. Scale bar, 5 μm. (B) A representative montage of time-lapse images showing dynamics of GFP-Mnr2 in a mature conidium. Arrowheads mark the GFP-Mnr2 puncta in all the three cells of the conidium. Numbers indicate time in minutes. Scale bar, 5 μm. (C) A montage of epifluorescence time-lapse images showing dynamic colocalization of GFP-Mnr2 puncta with a subpopulation of filamentous mitochondria stained with Mitotracker Deep Red FM. Arrowheads mark the GFP-Mnr2 puncta in close proximity of the mitochondria in old (conidium) cells. Numbers indicate time in seconds. Scale bar, 5 μm.

Membrane contact sites are dynamic in nature and respond to various metabolic cues. Given that Mnr2 in *S. cerevisiae* is a vCLAMP resident protein and that MCSs mediate transport of small molecules like lipids and ions, we hypothesized that vacuoles, as a reservoir of Mg^+2^, might come closer to mitochondria at the vCLAMPs to facilitate efficient uptake of Mg^+2^ ion. To test this hypothesis, we examined whether GFP-Mnr2 colocalized at all with the mitochondria during fungal development -more specifically, we observed the dynamics of both Mnr2 and mitochondria during appressorial development. Microscopic observations revealed that the GFP-Mnr2 puncta indeed colocalized with a subpopulation of mitochondria in the old conidial cells (Figure 3C). Further, such colocalizing GFP-Mnr2 puncta were dynamic and moved along and from one mitochondrial filament to another, likely at the vacuole-mitochondrion interface (Supplementary Movie 2). Importantly, such interactions between the vacuolar Mg^+2^ transporter and mitochondria were observed only in the older conidial cells, although the dynamic mitochondria were seen in both the developing young appressoria and germinating old conidial cells. Our observations suggest that Mnr2 is likely a vCLAMP member and helps in inter-organellar communication in *M. oryzae.*

### Mnr2-based Mg^+2^ homeostasis is required for mitochondrial integrity and function in aged fungal cells

Mitochondrial dysfunction is a hallmark of cellular aging. Having shown mitochondrial proximal localization of GFP-Mnr2 in the aged cells, we next examined whether the morphology and/or dynamics of the organelle were affected in the *mnr2*Δ mutant. In this direction, we studied the mitochondrial membrane potential (*ΔΨ*), as a readout for mitochondrial function using a Δ*Ψ*-dependent mitochondria-specific fluorescent dye 3, 3’-dihexyloxacarbocyanine iodide (DiOC_6_) (Pringle *et al.* 1989). While functional mitochondria in metabolically active cells are stained by DiOC_6_, those with a decreased membrane potential show reduced or no staining with the dye. In an *in vitro* assay, conidia of the wild-type or *mnr2*Δ *M. oryzae* were inoculated in complete medium to allow vegetative hyphal growth and subsequently stained with DiOC_6_. We observed that the mitochondria from both young (close to the tip) and old (close to the conidia) cells of the wild-type strain were stained uniformly (Figure 4A). However, interestingly, DiOC_6_-stained mitochondria were observed primarily in the younger cells of the *mnr2*Δ strain (Figure 4A). The mitochondria of the older cells of the *mnr2*Δ mutant were largely unstained, indicating that the membrane potential of the organelle would have been impaired therein. By quantifying the number of events of individual vegetative hypha stained with DiOC_6_, we found that approximately 75 % of the *mnr2*Δ old cells had decreased mitochondrial potential as compared with only 25 % seen in the wild-type (Figure 4B; *P* = 0.05). Taking together, these results reinforce the fact that Mnr2 plays a critical role in maintaining the mitochondrial membrane potential and/ or function, especially in the aged cells of *M. oryzae*.

**Figure 4:**
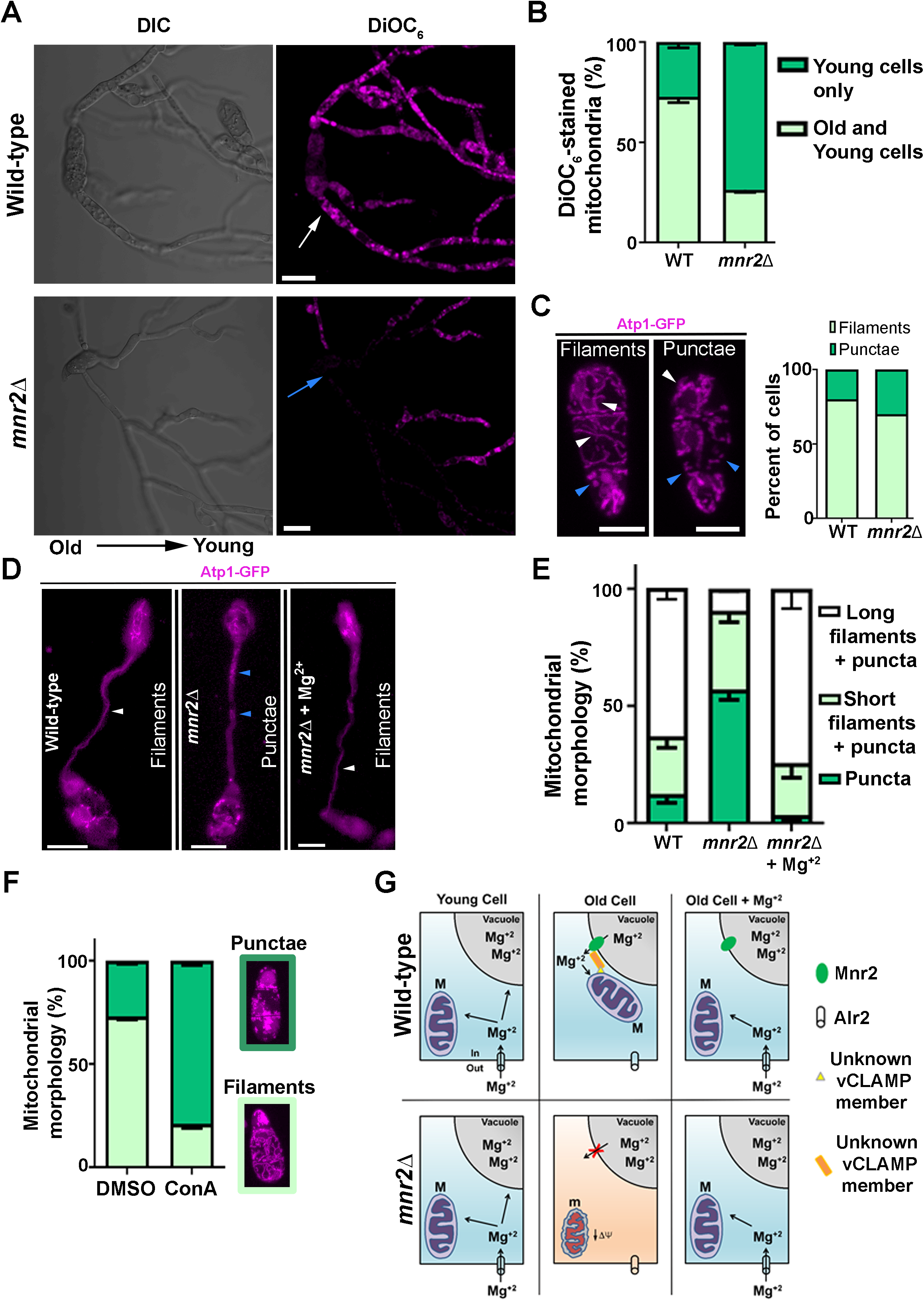
Mg^+2^ transporter Mnr2 is required for mitochondrial integrity and function in chronologically aged cells of *M. oryzae.* (A) Laser-scanning confocal micrographs depicting wild-type and *mnr2*Δ vegetative hyphae stained with DiOC_6_ to aid visualization of mitochondria with active membrane potential. The fluorescence images shown are the final z-projected images of the collected z-stacks. The white and blue arrows mark the DiOC_6_-stained and unstained mitochondria, respectively, especially in the aged hyphal cells. Scale bar, 10 μm. (B) Bar graph showing percentage of fungal microcultures containing mitochondria with active membrane potential, DiOC_6_-stained, in young and aged cells. Error bars show mean ± standard error of mean (SEM) from three replicates. n = 8 to 15 (wild-type) and 7 to 9 (*mnr2*Δ) microcultures per replicate. (C) Laser-scanning confocal micrographs showing mitochondrial morphology in the Atp1-GFP-expressing wild-type or *mnr2*Δ conidia. White and blue arrowheads mark the tubular and fragmented/ punctate mitochondria, respectively. The fluorescence images shown are the final z-projected images. Scale bar, 5 μm. Bar graph showing percent of cells with filamentous or punctate mitochondria. n = 50 (wild-type) and 60 (*mnr2*Δ). (D) Epifluorescence micrographs showing mitochondrial morphology in the wild-type or *mnr2*Δ *M. oryzae*, with or without exogenously added Mg^+2^, during pathogenic (appressorial) development at 6 hpi on a hydrophobic surface. White and blue arrowheads mark the tubular and fragmented mitochondria, respectively. Scale bar, 10 μm. (E) Bar graph depicting the quantitative analysis of the mitochondrial morphology in the strains and conditions mentioned in (D). The data represents mean ± SEM from three replicates. n = 20 to 61 (wild-type), 20 to 68 (*mnr2*Δ), and 20 to 95 (*mnr2*Δ+Mg^+2^) germinated conidia per replicate. (F) A bar graph depicting the effect of concanamycin A, a specific inhibitor of V-ATPase, on mitochondrial morphology in the wild-type conidia expressing Atp1-GFP. Data represents mean ± SEM from three replicates. n = 36 to 38 (solvent control – DMSO-treated) and 29 to 37 (ConcA-treated) conidia per replicate. (G) A model proposed to show how Mnr2 likely maintains the cytosolic pool of Mg^+2^ ion required for proper functioning of the mitochondria in the old cells. While Alr2 transports Mg^+2^ ions from the extracellular milieu to the cytoplasm, excess ions are stored in the vacuoles under nutrient-rich condition. However, ion homeostasis, during aging or nutrient-deplete conditions, is ensured by Mnr2-mediated access to the vacuolar pool of Mg^+2^ ions. Loss of the Mnr2 function affects Mg^+2^ homeostasis, leading to reduced mitochondrial membrane potential (*ΔΨ*) and integrity, which in turn triggers an early onset of chronological aging in *M. oryzae.*

Next, we studied the loss of integrity and dynamics of mitochondria in the absence of Mnr2 function, if any, in the Atp1-GFP/ *mnr2*Δ strain. We observed a modest 10% increase of unusually fragmented i.e. short filamentous and/ or punctate mitochondrial structures in the freshly harvested *mnr2*Δ conidia when compared to those of the wild-type (Figure 4C). When such conidia were inoculated on a hydrophobic surface for germination and subsequent appressorial development, the *mnr2*Δ mutant showed fragmented mitochondria in almost 90% of the germ tubes and conidial cells (old cells; Figure 4D & 4E; *P* = 0.05; Supplementary Movie 3). However, only ~35% of the wild-type germ tubes and conidial cells showed punctate or short filaments of the mitochondria (Figure 4E; Supplementary Movie 4). Further, the fragmented mitochondria in the *mnr2*Δ mutant displayed localized movement, whereas the predominantly long filamentous mitochondria of the wild-type were highly dynamic, with consistent fusion and fission of the organelle (Supplementary Movies 5 & 6). Strikingly, *mnr2*Δ mutant showed wild-type-like mitochondrial morphology and distribution upon exogenous supply of Mg^+2^ (Figure 4D & 4E; *P* = 0.05). Interestingly, not only the morphology but also the dynamics of the *mnr2*Δ mitochondria was restored upon exogenous supplement of Mg^+2^ (Supplementary Movie 7).

Subsequently, we sought to understand whether or not the loss of mitochondrial integrity and function in the *mnr2*Δ was indeed associated with altered vacuolar pH in the *mnr2*Δ mutant. To answer this, we treated the wild-type (Atp1-GFP) conidia with Concanamycin A, a specific inhibitor of V-ATPase (Droese *et al.* 1993), and observed its effect on the mitochondrial morphology. Indeed, the Concanamycin A-treated wild-type conidia, with likely altered vacuolar pH upon inhibition of the V-ATPase function, exhibited significantly higher number of punctate mitochondria when compared with the solvent control (Figure 4F; *P* = 0.05).

Given that the exogenous Mg^+2^ supplement prevented early cell death in the older part of the colony and restored the mitochondrial integrity and dynamics in the *mnr2*Δ mutant, we posit that Mnr2 provides access to the vacuolar pool of Mg^+2^ ion to the mitochondria likely at the MCSs, especially during starvation or old age (Figure 4G). Taken together, the vacuolar ion transporter Mnr2 maintains the function of mitochondria, and therefore survival fitness, especially during chronological aging in *Magnaporthe oryzae*.

## Discussion

In this study, we characterized a vacuolar CorA Mg^+2^ transporter Mnr2, a previously unknown regulator of aging, in *M. oryzae.* The *mnr2*Δ null mutant displayed loss in cell viability of the chronologically aged part of the colony which was rescued by exogenous Mg^+2^ supplement. Mnr2 localized as puncta onto the vacuolar membrane and displayed temporal expression, being higher in aged cells when compared with young cells. Our results demonstrate that Mnr2 contributes to the maintenance of vacuolar pH and canonical vacuolar function. This is evidenced by fragmented vacuoles, a hallmark of aging cells, along with *vma*-like phenotype (Kane 2006), with increased sensitivity to oxidative stress and alkaline pH, quinacrine-unstained vacuoles and decreased mitophagy in the *mnr2*Δ null mutant. Colocalization studies revealed that Mnr2 comes in close proximity with the mitochondria in aged cells. Cells lacking Mnr2 displayed a significant loss of tubular mitochondria during appressorium development and loss of mitochondrial function in the aged cells. Intriguingly, exogenous Mg^+2^ restored both the long-tubular mitochondrial structure and dynamics in cells lacking Mnr2. Taken together, our results demonstrate that the Mnr2 is essential for Mg^+2^ homeostasis and is important for the mitochondrial integrity and organellar cross-talk during chronological aging in the rice-blast fungal pathogen *Magnaporthe oryzae.*

Localization of a protein to spatially confined focal structures is commonly observed in the case of membrane contact sites like ERMES, vCLAMP, and NVJ (Eisenberg-Bord *et al.* 2016). The punctate GFP-Mnr2 localization as distinct patches onto the vacuolar membrane in *M. oryzae* was in line with the localization of Mnr2 in budding yeast (Elbaz-Alon *et al.* 2014), indicating a conserved subcellular localization in the two ascomycetes. Membrane contact sites have been shown to be dynamic and responsive to metabolic cues. The plasma membrane-ER contact site is induced upon Ca^+2^ store depletion (Wu *et al.* 2006), while NVJ shows enlargement as budding yeast enters stationary phase (Pan *et al.* 2000). *S. cerevisiae* shows increased vCLAMP-mediated contacts depending upon the carbon source within the cell (Honscher *et al.* 2014) and upon loss of the ERMES pathway (Elbaz-Alon *et al.* 2014). We also observed an increased expression of GFP-Mnr2 with an increase in the chronological age in *M. oryzae.* Regulators of lifespan are known to show differential expression and/ or distribution during the process of aging in *S. cerevisiae.* Protein levels of Sir2, an essential modulator of replicative lifespan, decline as cells age in *S. cerevisiae* (Dang *et al.* 2009). Similarly, the plasma membrane proton ATPase, Pma1, is expressed asymmetrically and accumulates in mother cells of *S. cerevisiae* (Henderson *et al.* 2014). Our results establish the link between Mnr2 expression and chronological age, which is also associated with nutrient depletion. It would be worth addressing whether Mnr2 could sense nutritional cues, likely via Target-of-Rapamycin (TOR) signaling cascade (Loewith and Hall 2011), and therefore be used as a potentially novel marker to delineate aged fungal cells.

Vacuoles are dynamically regulated both in number and size and respond to various environmental and physiological conditions including aging. Overexpression of Osh6 in *S. cerevisiae* has been shown to enhance vacuolar fusion leading to an increase in replicative lifespan (Gebre *et al.* 2012). The *mnr2*Δ mutant cells, with a decreased lifespan, showed small spherical vacuoles in a significantly increased number of conidial cells. Acidic pH of the vacuole, regulated by the V-ATPase, is central to its function of ion homeostasis and being storage reservoir for metal ions, amino acids, carbohydrates, and other metabolites (Kane 2006). A recent study on Sch9 kinase, a resident of vacuolar membrane in *S. cerevisiae*, also links TOR signaling to V-ATPase function regulating vacuolar acidity and cellular lifespan (Wilms *et al.* 2017). Mnr2, by exporting Mg^2+^ from vacuoles to cytosol, plays a role in maintaining the vacuolar Mg^2+^ levels and pH likely in fine co-ordination with i) Mg^2+^/H^+^ exchanger; importing Mg^2+^ from cytosol to vacuoles while pumping out H^+^ and ii) V-ATPase; importing H^+^ within the vacuoles in *S. cerevisiae* (Pisat *et al.* 2009). The *Magnaporthe mnr2*Δ mutant displayed *vma*-like phenotype (Kane 2006), with growth defects in the presence of oxidative stress and at alkaline pH. Vacuolar acidification is intimately linked with autophagy induction and *vma* mutant also shows a defect in protein degradation induced by starvation (Nakamura *et al.* 1997). The *mnr2*Δ mutant in our study also showed significantly lower number of quinacrine-stained vacuoles upon nutrient-starvation conditions, and as a consequence, a lower autophagic turnover of mitochondria. In summary, our results suggest that the vacuolar Mg^+2^ transporter, through a fine balance of vacuolar pH, prevents age-associated decline in the vacuolar function. However, it remains to be seen if the expression of Mnr2 is dependent on the vacuolar pH or *vice versa* in *M. oryzae.*

Organelles within a cell come in close proximity at MCSs to exchange metabolites, ions, and lipids (Elbaz-Alon *et al.* 2014). One of the striking observations of this study was that the GFP-Mnr2 puncta were dynamic and moved along the vacuolar periphery, and colocalized with mitochondria in the chronologically aged cells in *M. oryzae.* To our knowledge, such dynamic behavior for any resident proteins of the MCSs has not been reported thus far. A functional significance of colocalization of Mnr2 with mitochondria remained elusive (Elbaz-Alon *et al.* 2014). A previous study in *S. cerevisiae* identified Avt1, a neutral amino acid transporter mediating proton-dependent neutral amino acid storage in the vacuoles, necessary for the cross-talk between vacuolar pH and mitochondrial function and therefore lifespan (Hughes and Gottschling 2012). Our findings show that the Mnr2 function is required for mitochondrial integrity and function, especially in the aged cells of *M. oryzae*. Significantly, a higher proportion of *mnr2*Δ cells with fragmented mitochondria during appressorium formation is in agreement with our earlier observation of the increased demand of Mg^+2^ during pathogenic development (Reza *et al.* 2016). Importantly, this study uncovers a novel, Mnr2-mediated organellar cross-talk between vacuoles and mitochondria, especially during aging in a filamentous fungus.

We propose that the damaged mitochondria with sub-optimal function accumulate with age and require access to Mg^+2^ ions either available in the cytoplasm or previously stored in the vacuole for normal function. In the absence of Mnr2 function, vacuolar Mg^+2^ ions would be inaccessible, and hence an Alr2-based replenishment of cytoplasmic pool of Mg^+2^ ion should be able to support mitochondrial integrity and function. Indeed, exogenous Mg^+2^ supplements restored the integrity and kinetics of mitochondrial fission-fusion in *mnr2*Δ mutant, by likely replenishing the cytosolic pool of Mg^2+^ ions and making it available for mitochondrial uptake. Collectively, our findings highlight a crucial role of the vacuolar transporter Mnr2 in providing access to the stored Mg^+2^ ions at the vacuole-mitochondria interface, to maintain the mitochondrial integrity and cellular energy status, especially in the chronologically older cells, increasing survival fitness during aging in *M. oryzae* (Figure 4G).

The blast fungus *Magnaporthe oryzae*, causing devastating rice and wheat losses (Wilson and Talbot 2009; Islam *et al.* 2016), tops the list of plant pathogens owing to its economic significance (Dean *et al.* 2012) and has emerged as a model system for the study of host-pathogen interactions. Several studies have highlighted the significance of mitochondria in aging in different model organisms, while others have dissected the role of vacuoles towards longevity. Furthermore, studies highlighting the role of inter organellar crosstalk in mediating lifespan extension in various model organisms is limited. Importantly, the mechanistic details regulating lifespan in *M. oryzae*, an important plant pathogen, have been undervalued. This study on Mnr2-mediated protection of mitochondrial dysfunction demonstrates a novel strategy of safeguarding cells in aged cells. How Mnr2 senses chronological age of a cell and triggers its heightened expression remains to be addressed.

## Experimental procedures

### Fungal strains, culture conditions, and transformation

*Magnaporthe oryzae* B157 strain (MTCC accession number 12236), belonging to the international race IC9, was previously isolated in our laboratory in collaboration with Indian Institute of Rice Research, Hyderabad (Kachroo *et al.* 1994). The fungus was grown and maintained on prune agar (PA). Complete medium (CM) was used for growing fungal biomass for DNA isolation and microscopy.

Vegetative growth was measured in terms of colony diameter on yeast extract dextrose (YEGA) agar. Conidia were harvested after growing the cultures on PA for 2 - 3 days in dark, followed by growth under constant illumination for the next 7 - 8 days and total conidia were harvested as described earlier (Patkar *et al.* 2012).

Gene-tagging and deletion constructs were transferred into *M. oryzae* either by protoplast transformation or *Agrobacterium tumefaciens-mediated* transformation (Mullins *et al.* 2001; Patkar *et al.* 2015). The transformants were selected on YEGA with 300 μg ml^-1^ Zeocin or Basal medium with 100 μg ml^-1^ Chlorimuron ethyl or 50 μg ml^-1^ Glufosinate Ammonium. The transformants were screened by locus-specific PCR and further validated by Southern blot hybridization.

For the growth assay on YEGA with different pH values and in presence of H_2_O_2_ (5 mM and 6 mM), the wild-type and *mnr2*Δ strains were inoculated and imaged after 5 dpi and 10 dpi respectively.

### Plasmid constructs

Mnr2 was tagged at the N-terminus with GFP using marker-fusion tagging (Lai *et al.* 2010) by targeted replacement of the native orf with *BAR-GFP-MNR2* sequence. Here, first 1195 bp *MNR2* orf was amplified using primers Mnr2 F_KpnI/Mnr2 R_Hindlll and then cloned in pFGL718 in-frame with *BAR-GFP* to get pRPL045. The *MNR2* promoter region was digested out from pBSKS-*MNR2* promoter using Sall and Spel and cloned at the same sites in pRPL045 to get pRPL046. All the constructs were confirmed by restriction enzyme digestion. All the PCR were carried out using proof-reading XT-5 polymerase enzyme (GeNei). The construct for tagging of Mnr2 was first transferred into *Agrobacterium tumefaciens* by electroporation and then to *M. oryzae* strain B157 via ATMT method. The tagged strain was screened by PCR, validated by Southern blot hybridization, and analyzed by fluorescence microscopy.

Mnr2_Nest_F and Venus_R_Kpnl (with a stop codon) were used to amplify *MNR2 promoter-BAR-GFP* (2181 bp) from pRPL046, with a stop codon at the end of GFP-coding frame, to drive cytosolic expression of the reporter gene by *MNR2* promoter. The PCR product was used to transform protoplasts of wild-type strain, B157. The transformants were selected by growth on selection medium and then by microscopy and PCR.

For *MNR2* deletion in Atp1-GFP-expressing strain, a 3053 bp *MNR2*-deletion cassette was PCR-amplified from *mnr2*Δ genomic DNA, using Mnr2_Nest_F/Mnr2_Nest_R, and was transferred in Atp1-GFP protoplasts using the previously described method. Transformants with a targeted gene deletion were screened by locus-specific PCR and validated by Southern blot hybridization.

### Staining protocols

Vacuoles were visualized by staining with 7-amino-4-Chloromethylcoumarin (CMAC) (ThermoFischer Scientific, Cat. No. C2110; CellTracker^TM^ Blue CMAC dye) dissolved in DMSO to give a 10 mM stock. Briefly, the harvested conidia, germinating conidia, or appressoria were incubated with 10 μM dye at 37 °C for 1 h 30 min. The cells were washed with 1x PBS thrice with 5 min incubation each and were imaged.

For phloxine B staining assay, the wild-type and *mnr2*Δ strains were cultured on YEGA containing 10 μM final concentration of the vital dye and imaged on 14 dpi for the overall lifespan assay or 7 dpi for the amiloride HCl treatment assay.

To visualize mitochondrial potential, 3, 3’-dihexyloxacarbocyanine iodide (DiOC_6_) (ThermoFischer Scientific; Cat. No. D273) staining was performed. Briefly, the wild-type and *mnr2*Δ conidia were allowed to germinate and form hyphae overnight in liquid complete medium. The germinated spores and vegetative hyphae were gently washed with solution containing 10 mM HEPES, pH 7.6 and 5% glucose. The cells were resuspended in the same washing solution containing 60 nM DiOC_6_ and incubated at room temperature for 1 h 30 min. The cells were washed twice with 10 mM HEPES, pH 7.6 and 5% glucose and resuspended in the same buffer for imaging.

For mitochondrial staining, the fungal samples were incubated in complete medium for appropriate time and then stained by adding 100 nM MitoTracker^TM^ Deep Red FM (ThermoFischer Scientific; Cat. No. M22426) to the culture for 30 min. The samples were imaged after briefly washing away the excess dye.

FM4-64 staining was performed to visualize the vacuolar membrane. Briefly, after 2 h of conidial germination, FM4-64 (ThermoFischer Scientific; Cat. No. T3166) was added at a concentration of 7.5 μM for 1 h. The conidia were gently washed twice and further incubated for 1 h before microscopy.

To assess the pH of the vacuolar lumen, both the wild-type and *mnr2*Δ strains were grown in complete medium for 48 h, and then transferred to water for 3 h before staining with 1 μg ml^-1^ quinacrine for 15 min.

### Microscopy and image processing

Subcellular localization was studied by laser-scanning confocal microscopy using an LSM 700/LSM 880 inverted confocal microscope (Carl Zeiss Inc., Germany), equipped with a Plan-Apochromat 63x/1.40 oil immersion lens. Imaging of eGFP and DiOC_6_-stained cells was carried out at 488/509 nm excitation/emission wavelengths. The CMAC-stained cells were imaged at 353/466 nm excitation/emission wavelengths. For live-cell imaging, fungal cultures were inoculated on glass-bottom petri dishes. The maximum projection images were obtained from Z stacks of 0.5 μm-spaced sections and processed and analyzed using ImageJ (https://imagej.nih.gov/ij/download.html) and/or Adobe Photoshop CC 2018 software.

### Analyses and quantification

Quantification of GFP-Mnr2 signal intensity was carried out after the projection of Z-stacks and was represented as the corrected total cell florescence (CTCF). Briefly, after Z projection, while GFP-Mnr2 signal was measured from the puncta, the background noise was calculated from a region surrounding the GFP-Mnr2 puncta, and the CTCF values were calculated as: Integrated density - (Area of selected region X mean fluorescence of background readings) and plotted using GraphPad Prism 6.00.

### Statistical analyses

Statistical analyses were done using GraphPad Prism 6.0 software. Student’s *t*-test followed by Unpaired *t*-test for comparing two groups and one-way ANOVA followed by Dunnett’s multiple comparison test for multiple groups were used and has been mentioned in the figure legends where applicable. The data sets represented as bar graphs were analyzed using Two-way ANOVA followed by Tukey’s multiple comparison test.

### Drugs used

Concanamycin A (Sigma Aldrich; Cat. No. C9705) was used as a specific inhibitor of V-ATPase. Wild-type conidia were treated with Concanamycin A at a concentration of 500 nM for 6 h to study the effect of loss of V-ATPase function on mitochondrial integrity.

Amiloride HCl (Sigma Aldrich; Cat. No. A7410) was used as a specific inhibitor of Mg^+2^/ H^+^ exchanger. Wild-type and *mnr2*Δ cultures were inoculated on YEGA supplemented with 1 mM Amiloride HCl. A no-drug sample with 1% DMSO was used as a solvent control in the assay. The cultures were imaged on 7 dpi.

### Mitophagy assay

Mitophagy assay was performed as explained previously (He *et al.* 2013) with a few modifications. Briefly, Atp1-GFP/ wild-type and *mnr2*Δ/ Atp1-GFP strains were grown in complete medium (CM) containing 1% Glucose for 48 h. The cultures were gently washed thrice with milliQ water before replacing CM with 1% Glucose with CM-glucose + 1.5% (v/v) glycerol (to induce mitochondrial biogenesis) and the germinated conidia were grown for another 33 h. The cultures were gently washed thrice with milliQ water and shifted to minimal media (MM) without NaNo_3_ (nitrogen starvation to induce mitophagy) + 1% glucose for 7 h. Wild-type with 3 mM PMSF was used as a negative control for mitophagy. The cells were stained with CMAC (as explained earlier) to mark the vacuoles and imaged at appropriate time.

## Acknowledgements

We thank N. Naqvi (TLL, Singapore) for sharing reagents. MHR acknowledges late B.B. Chattoo and late J. Manjrekar for infrastructural and financial support during initial part of this work at M S University, Vadodara. We also acknowledge H. Shah for her assistance in strain construction, critical reading of the manuscript, and valuable suggestions. MHR received funding for this work as a National Postdoctoral Fellow (PDF/2016/002858) from Science and Engineering Research Board (SERB), Department of Science and Technology (DST), Government of India and is currently a Department of Biotechnology-Research Associate-I of Government of India. The award of Tata Innovation Fellowship (BT/HRD/35/01/03/2017) and JNCASR intramural funding support to KS is acknowledged. This work was supported by the Ramalingaswami Fellowship, DBT, Government of India (BT/RLF/Re-entry/32/2014) awarded to RP. This work was also supported by Department of Biotechnology (DBT) grant in Life Science Research, Education and Training at JNCASR (BT/INF/22/SP27679/2018). We thank the members of the Bharat Chattoo Genome Research Centre Group and Molecular Mycology Laboratory for valuable discussions.

## Conflict of Interest

The authors declare that they have no conflict of interest.

